# Recruiting ESCRT to single-chain heterotrimer peptide-MHCI releases antigen-presenting vesicles that stimulate T cells selectively

**DOI:** 10.1101/2025.01.17.633600

**Authors:** Blade A. Olson, Kathryn E. Huey-Tubman, Zhiyuan Mao, Magnus A. G. Hoffmann, Richard M. Murray, Stephen L. Mayo

## Abstract

Immune cells naturally secrete extracellular antigen-presenting vesicles (APVs) displaying peptide:MHC complexes to facilitate the initiation, expansion, maintenance, or silencing of immune responses. Previous work has sought to manufacture and purify these vesicles for cell-free immunotherapies. In this study, APV assembly and release is achieved in non-immune cells by transfecting HEK293T or Expi293F cells with a single-chain heterotrimer (SCT) peptide/major histocompatibility complex I (pMHCI) construct containing an ESCRT- and ALIX-binding region (EABR) sequence appended to the cytoplasmic tail; this EABR sequence recruits ESCRT proteins to induce the budding of APVs displaying SCT pMHCI. A comparison of multiple pMHCI constructs shows that inducing the release of APVs by the addition of an EABR sequence generalizes across SCT pMHCI constructs. Purified pMHCI/EABR APVs selectively stimulate IFN-γ release from T cells presenting their cognate T cell receptor, demonstrating the potential use of these vesicles as a form of cell-free immunotherapy.

**Significance Statement:** Immune cells are known to naturally release pMHC-displaying extracellular vesicles (EVs), called antigen-presenting vesicles (APVs), which can orchestrate immune responses either directly or with the aid of antigen-presenting cells (APCs). For decades, researchers have pursued ways to replicate these APVs for immunotherapy by using chemically modified nanoparticles or by engineering the increased expression of APVs from immune cells which are typically low yield. Here we presents a broadly applicable platform for generating high concentrations of pMHCI-displaying APVs that can selectively modulate T cells, demonstrating a significant advance in the engineering of APVs for cell-free immunotherapy. The APVs presented here, and related APVs, could be translated into clinical therapies for modulating cancer progression or regulating autoimmunity in addition to their use as a tool to help characterize how endogenous extracellular vesicles influence the immune system.

## Introduction

Almost thirty years ago, Raposo, G. et al. first reported on the release of B cell-derived extracellular vesicles (EVs) with functional peptide:MHC (pMHC) complexes on their surface.^1^ Since then, EVs that were previously labeled as simple cellular “debris” have been increasingly recognized for their immunoregulatory potential as a simpler, cell-free alternative to cell-based immunotherapies.^2,3^ Naturally occurring pMHC-displaying antigen-presenting vesicles (APVs) in particular have been demonstrated to orchestrate immune responses either directly or with the aid of antigen-presenting cells (APCs).^4^ Research teams have ventured to engineer more complex versions of these antigen-presenting vesicles: whole mammalian cells conjugated with immunostimulatory molecules,^5–9^ nanoparticles coated with APC-derived membranes via extrusion or sonication,^10,11^ or engineered extracellular blebs from dendritic cells.^12–16^ These previous approaches demonstrated the therapeutic potential of APVs, but production is often low yield or requires the patient’s own cells to be harvested and processed before treatment can be administered. Significant attention has been given to manufacturing APVs via microfluidics,^17^ but existing microfluidic vesicles suffer from short half-lives and are susceptible to rupture due to osmotic changes and the turbulent flow of the lymphatic and cardiovascular systems. Polymer-coated iron-oxide nanoparticles conjugated with pMHCI have shown remarkable therapeutic potential for autoimmune diseases such as type 1 diabetes, but the stiff spherical structure of the nanoparticle scaffold reduces the potential surface area for ideal pMHC:T cell receptor (TCR) interactions;^18–20^ additionally, pMHC complexes are rigidly adhered to the nanoparticle surface, limiting pMHC movement and clustering considered important for the formation of a T cell stimulating immunological synapse.^21^ Optimal, engineered APVs would ideally recreate the endogenous form of APVs, while maximizing the concentration of proteins that are critical for immunoregulation.

Here, we describe the engineered assembly and budding of peptide/MHCI-displaying APVs from the cell surface of non-immune cells by DNA transfection of an engineered single-chain heterotrimer (SCT) peptide/MHCI (pMHCI). Building on a recently described method for generating enveloped virus-like particles displaying the SARS-CoV-2 spike protein,^22^ we fused an endosomal sorting complex required for transport (ESCRT)- and ALG-2-interacting protein X (ALIX)-binding region (EABR) to the cytoplasmic tail of SCT pMHCI to directly recruit the ESCRT machinery and induce the release of pMHCI-displaying vesicles. Immuno-electron microscopy (immuno-EM) of the supernatant from transfected cells showed halos of black punctate surrounding exosome-like vesicles, confirming the presence of SCT pMHCI on APVs. ELISPOT assays show that SCT pMHCI/EABR APVs are capable of stimulating IFN-γ release from T cells at a level equivalent to refolded pMHCI tetramer, and that the APVs selectively stimulate T cells presenting their cognate TCR. Altogether, these results demonstrate a new method for generating synthetic, native-like APVs in high copy number and valency with the therapeutic potential for improved cell-free immunoregulation.

## Results

### EABR addition to the cytoplasmic tail of SCT pMHCI promotes APV release

To generate APVs from non-immune cells, we attached an EABR sequence from the human CEP55 protein^23^ to the cytoplasmic tail of MHCI (Fig. 1A). To ensure each APV presented the same, intended peptide on MHCI, the complete peptide:MHCI complex was formed into a single-chain heterotrimer; the N-terminus of the HLA-A*02:01 alpha chain was extended with a 4xG_4_S linker connecting to the C-terminus of beta-2 microglobulin (β2M), which was itself extended at its N-terminus by a 3xG_4_S linker connecting to either the 10-mer MART-1 (MART1) or 9-mer NY-ESO-1 (NYESO1) cancer-related peptides.^24^ The EABR sequence used here was identical to the sequence used by Hoffmann et al.^22^ to create enveloped virus-like particles (eVLPs) displaying the SARS-CoV-2 spike protein, and was attached to the cytoplasmic tail of the HLA-A*02:01 alpha chain by a G_3_S linker (Fig. 1B). Adopting the nomenclature of Chour et al.’s thermostability studies^25,26^ on previous SCT pMHCI designs, our first SCT pMHCI construct was named “D1/EABR” to denote a “D1” SCT design with an “EABR” sequence appended to the end of its cytoplasmic tail. A second “D9” variant of SCT pMHCI, containing Y84C and A139C substitutions, was also tested given its reported improvements in thermostability and T cell receptor binding compared to the D1 design.^27^

**Figure 1.**
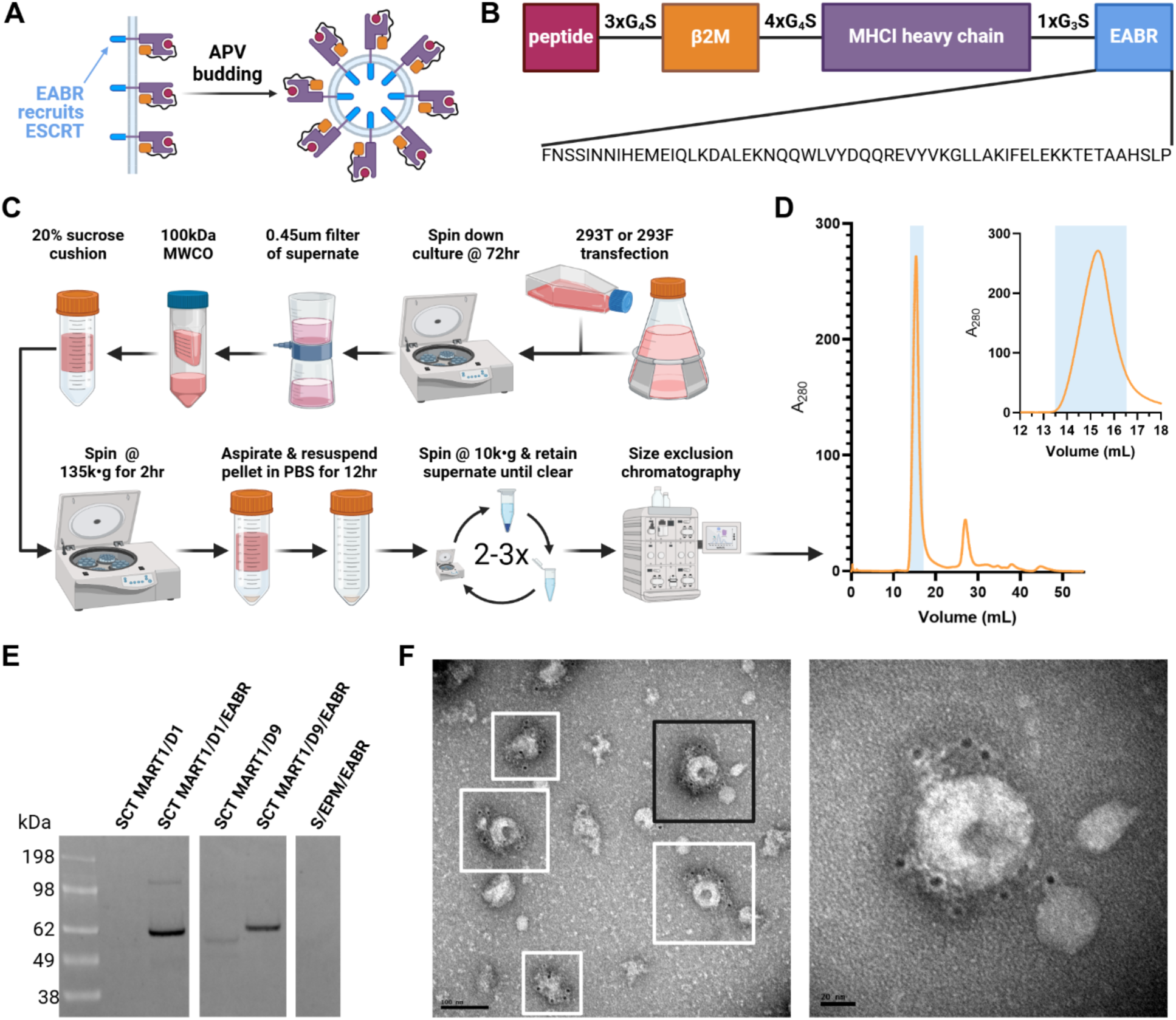
EABR addition to the cytoplasmic tail of SCT pMHCI promotes APV budding and release. (A) Schematic of membrane-bound SCT pMHCI proteins on a cell surface containing cytoplasmic tail EABR additions that induce budding of an APV comprising a lipid bilayer with embedded SCT pMHCI proteins. (B) SCT pMHCI/EABR construct. Top: the SCT pMHCI protein is composed of a peptide fused to a 3x(Gly_4_Ser) linker, β2M, a 4x(Gly_4_Ser) linker, HLA-A*02:01 alpha chain, a Gly_3_Ser spacer, and an EABR sequence. Bottom: EABR sequence. (C) Schematic showing production and purification of APVs. (D) Chromatogram representative of the typical yield of SCT pMHCI APVs from a 100 mL transfection of Expi293F cells. Inset highlights void volume peak. (E) Western blot analysis detecting SCT pMHCI protein from APVs purified by ultracentrifugation on a 20% sucrose cushion from transfected HEK293T cell culture supernatants. Cells were transfected with SCT MART1/D1, SCT MART1/D1/EABR, SCT MART1/D9, SCT MART1/D9/EABR, and SARS-CoV-2 spike(S)/EPM/EABR (S/EPM/EABR) constructs. Rabbit anti-HLA-A primary antibody and Alexa488-conjugated anti-rabbit secondary antibody were used to detect SCT pMHCI protein. (F) Immuno-EM images of SCT MART1/D1/EABR APVs purified from transfected Expi293F cell culture supernatants by ultracentrifugation and SEC. Primary rabbit anti-HLA-A antibody identical to primary antibody used in (E). Secondary 6 nm gold-conjugated anti-rabbit antibody appears as black punctate in image. Left: representative APVs are highlighted in boxes. Scale bar, 100 nm. Right: close up of black-boxed APV. Scale bar, 20 nm.

APVs were expressed by transfecting HEK293T or Expi293F cells with DNA plasmids encoding the SCT pMHCI constructs. Vesicles were purified by ultracentrifugation of cell culture supernatant on a 20% sucrose gradient as is typical for the purification of viral particles and exosomes^28^ (Fig. 1C). Ultracentrifuged product was resuspended in pH

7.4 phosphate-buffered saline (PBS) and optionally further purified by size exclusion chromatography (SEC) to yield a high concentration of product in the void volume (Fig. 1D). Western blot analysis of the purified product showed that the addition of the EABR sequence to the cytoplasmic tails of both the MART1/D1/EABR and MART1/D9/EABR constructs generated higher levels of MHCI compared to their non-EABR counterparts (Fig. 1E). Because pMHCI is difficult to resolve with cryo-EM or cryo-ET, we prepared immuno-electron microscopy grids of the purified product to confirm the presence of pMHCI-displaying APVs, which presented as cup-like, collapsed spheres resembling the canonical shape of exosomes that partially collapse upon paraformaldehyde fixation and negative staining^29^ (Fig. 1F and S1). Western blots of these pMHCI vesicles did not stain for the hallmark exosomal tetraspanin proteins CD81 or CD63 (Fig. S2); as a result, the vesicles have been denoted as pMHCI-displaying “antigen-presenting vesicles” to differentiate them from traditional “exosomes.”

### EABR addition to SCT pMHCI promotes budding of the complete, intended pMHCI complex on APVs

To better quantify the relative improvement in APV formation with EABR addition, we performed ELISA of SCT MART1 and SCT NYESO1 constructs. Both showed over 100-fold improvements in APV formation with the addition of the EABR sequence (Fig. 2A). Subsequent ELISAs showed that appending EABR to MHCI heavy chain (HC) alone (i.e., without covalently linked β2M and presenting peptide) still boosted MHCI yield in the purified transfection product relative to HC without an EABR tag, and an even greater concentration of MHCI resulted from transfection with EABR-tagged single-chain heterodimer (SCD) constructs comprised of MHCI heavy chain covalently linked to β2M without a presenting peptide (Fig. S3). Consequently, we next investigated whether the SCT form of pMHCI was necessary to assemble the intended pMHCI complex on released APVs.

**Figure 2.**
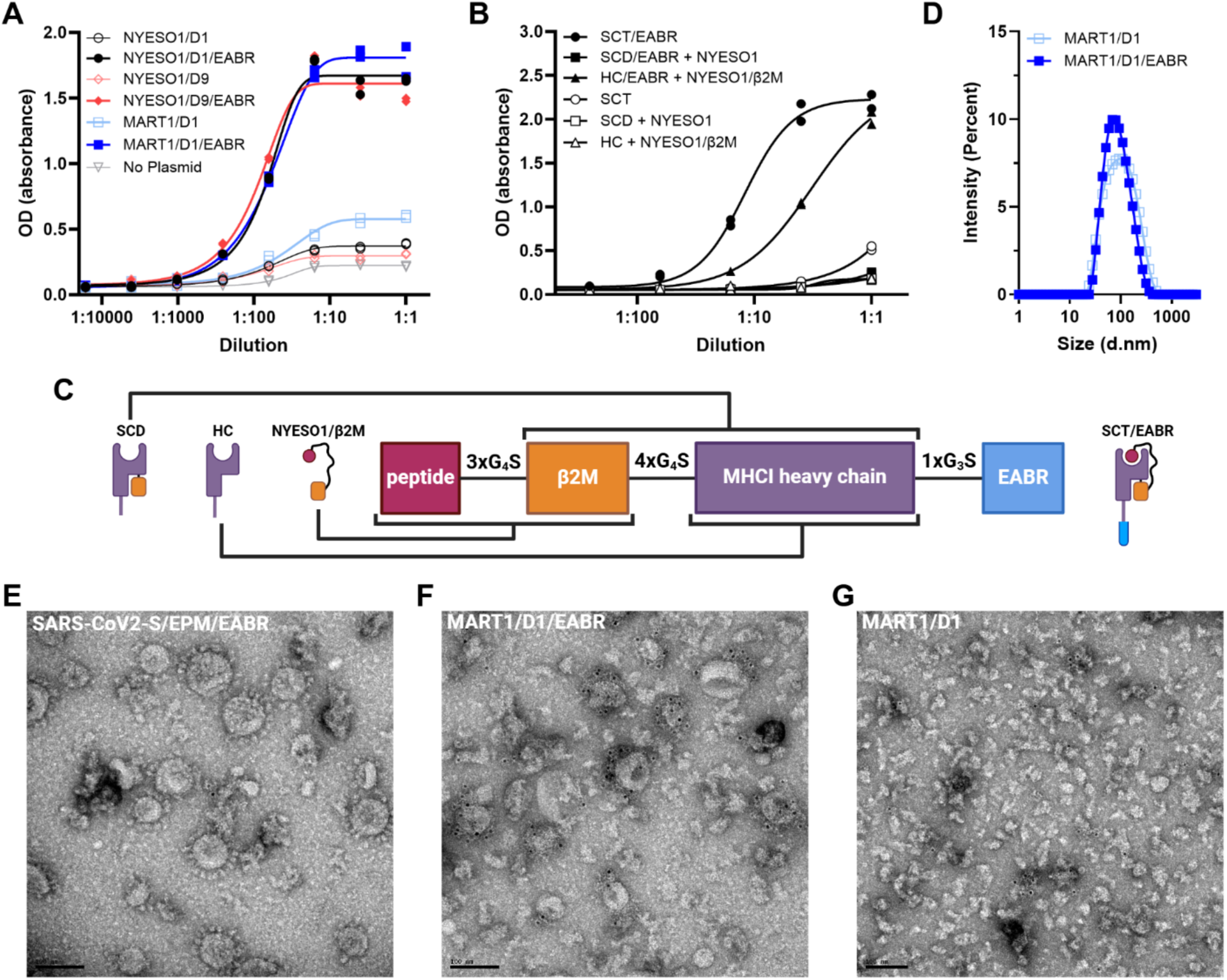
EABR addition to SCT pMHCI promotes budding of the complete, intended pMHCI complex on APVs. (A) Indirect ELISA of APVs purified by ultracentrifugation from transfected HEK293T cells with a primary rabbit anti-HLA-A antibody and secondary HRP-conjugated anti-rabbit antibody. Cells were transfected with SCT NYESO1/D1, SCT NYESO1/D1/EABR, SCT NYESO1/D9, SCT NYESO1/D9/EABR, SCT MART1/D1, and SCT MART1/D1/EABR constructs (2 replicates per dilution). (B) Sandwich ELISA of APVs purified by ultracentrifugation from transfected HEK293T cells using anti-NYESO1:HLA-A*02:01 (3M4E5) for capture, biotinylated 3M4E5 for detection, and HRP-conjugated streptavidin secondary. Cells were transfected with SCT NYESO1/D9/EABR, SCD D9/EABR and NYESO1 peptide (SCD/EABR + NYESO1)), HC D9/EABR and NYESO1/β2M (HC/EABR + NYESO1/β2M), SCT NYESO1/D9 (SCT), SCD D9 and NYESO1 peptide (SCD + NYESO1), HC D9 and NYESO1/β2M (HC + NYESO1/β2M) constructs (2 replicates per dilution). (C) Schematic showing full SCT pMHCI/EABR construct (SCT/EABR) and various alternative constructs used for the ELISA in (B): NYESO1/β2M, HC, and SCD. (D) Dynamic light scattering measurement of SCT MART1/D1 and SCT MART1/D1/EABR APVs represented as a probability mass function of the expected diameter values for each sample. Points are connected by a simple interpolating spline. (E-G) Immuno-EM micrographs of different EABR vesicles purified from transfected Expi293F cell culture supernatants by ultracentrifugation and SEC. Samples were stained with a primary rabbit anti-HLA-A antibody and a secondary 6 nm gold-conjugated anti-rabbit antibody that appears as black punctate in image. Scale bars, 100 nm. (E) SARS-CoV2-S/EPM/EABR. Spike protein coronas are visible around the edges of the spherical vesicles. A single gold nanoparticle from a secondary antibody is visible in the middle of the image. (F) SCT MART1/D1/EABR gold particles from secondary antibody can be seen as halos of black punctate around deflated vesicles. (G) SCT MART1/D1 scattered gold particles from secondary antibody can be seen as non-specific staining without the presence of vesicles.

To compare the impact of different NYESO1/MHCI construct designs on APV assembly and release, we selectively removed the Gly_4_Ser linkers from the SCT NYESO1/D9 construct – either removing the 3xG_4_S linker between the peptide and β2M yielding SCD, or the 4xG_4_S linker between β2M and the MHCI heavy chain yielding NYESO1/β2M – and performed ELISAs with a TCR-like antibody specific for the fully formed NYESO1:HLA-A*02:01 complex (Fig. 2B & 2C).^30^ Decoupling the NYESO1 peptide from the EABR-tagged SCD MHCI complex led to an over 10-fold reduction in the detected level of expression of NYESO1/MHCI APVs in HEK293T cells (Fig. 2B). Co-transfecting two separate constructs of HC/EABR and NYESO1/β2M generated APVs detectable by sandwich ELISA and provides an alternative design for APVs; however, the SCT form of NYESO1/D9/EABR provided the best expression of pMHCI-displaying APVs out of all constructs tested (Fig. 2B), and it is the only design that ensures the APVs are presenting the construct’s intended peptide.

Interestingly, we still recovered low levels of MHCI protein after purification of non-EABR SCT pMHCI, an effect that was amplified by expressing the same constructs in the higher-expressing Expi293F cell line, which has been used previously for exosome production^31^ (Fig. 2A & S4). Dynamic light scattering (DLS) of both the purified SCT MART1/D1 and SCT MART1/D1/EABR constructs showed similar median particle diameters of 60-80 nm (range 30-250 nm) with the SCT MART1/D1/EABR construct exhibiting a slightly narrower distribution compared to the non-EABR SCT MART1/D1 construct (Fig. 2D). This particle size distribution is consistent with the vesicle sizes seen in immuno-EM images of the EABR constructs. We were unable, however, to observe pMHCI vesicles when imaging immuno-EM grid preparations of the SCT MART1/D1 construct purified from Expi293F supernatant. Instead, the non-EABR SCT MART1/D1 grids showed what appeared to be pMHC aggregates or blebbing of sparsely populated membrane, which was less frequently observed in the grids prepared for the EABR-tagged vesicles (Fig. 2F, 2G, & S1). As a direct contrast, identically prepared EM grids for both SARS-CoV-2 S/EPM/EABR and MART1/D1/EABR constructs were densely populated with vesicles displaying their respective proteins, and only the MART1/D1/EABR vesicles stained positively with a halo of anti-HLA-A antibody around each vesicle (Fig. 2E & 2F). Altogether, our results show that SCT pMHCI promotes the display of complete pMHCI complexes on APVs, and that addition of the EABR sequence increases APV release.

### EABR-mediated APV release generalizes across SCT pMHCI variants

To further explore the generalizability of EABR sequence addition and heterotrimerization of pMHCI on APV release, we tested whether EABR-tagged SCT constructs of murine MHCI and chimeric human/mouse MHCI would also generate APVs displaying pMHCI relevant for in vivo studies of immunoregulation. SCT constructs of murine MHCI H-2Kd were prepared as described for the human D1 constructs with either an NRP-V7 or IGRP_206-214_ type 1 diabetes-relevant presenting peptide fused to a 3x(Gly_4_Ser) linker, murine β2M, a 4x(Gly_4_Ser) linker, and the H-2Kd alpha chain, i.e., NRP-V7/H-2Kd and IGRP_206-214_/H-2Kd, respectively. The chimeric human/mouse MHCI HLA-A2/H-2Db and HLA-A2/H-2Kb were designed according to the D9 SCT design, with an NYESO1 presenting peptide fused to a 3x(Gly_4_Ser) linker, human β2M, a 4x(Gly_4_Ser) linker, and a chimeric MHCI alpha chain composed of the extracellular α1 and α2 domains of human Y84A,A139C HLA-A*02:01 heavy chain and the α3, transmembrane, and cytoplasmic domains of murine H-2Kb or H-2Db heavy chain, i.e., NYESO1/A2_D9/H-2Db and NYESO1/A2_D9/H-2Kb, respectively.

For both SCT NRP-V7/H-2Kd and SCT IGRP_206-214_-V7/H-2Kd, addition of an EABR sequence to the cytoplasmic tail of MHCI heavy chain amplified the production of APVs as detected by sandwich ELISA (Fig. 3A). However, like the previous SCT MART1/D1, NYESO1/D1, and NYESO1/D9 pMHCI constructs, the SCT NRP-V7/H-2Kd pMHCI construct showed evidence of substantial vesicle production in HEK293T cells regardless of the addition of the EABR sequence. Interestingly, IGRP_206-214_-V7/H-2Kd vesicle production was noticeably reduced despite the only sequence difference being a change in the presenting peptide; appending an EABR sequence to this IGRP_206-214_-V7/H-2Kd construct led to a more dramatic change in APV production relative to EABR’s effect on SCT NRP-V7/H-2Kd pMHCI APV production. Nonetheless, addition of an EABR sequence for both constructs amplified vesicle production.

**Figure 3.**
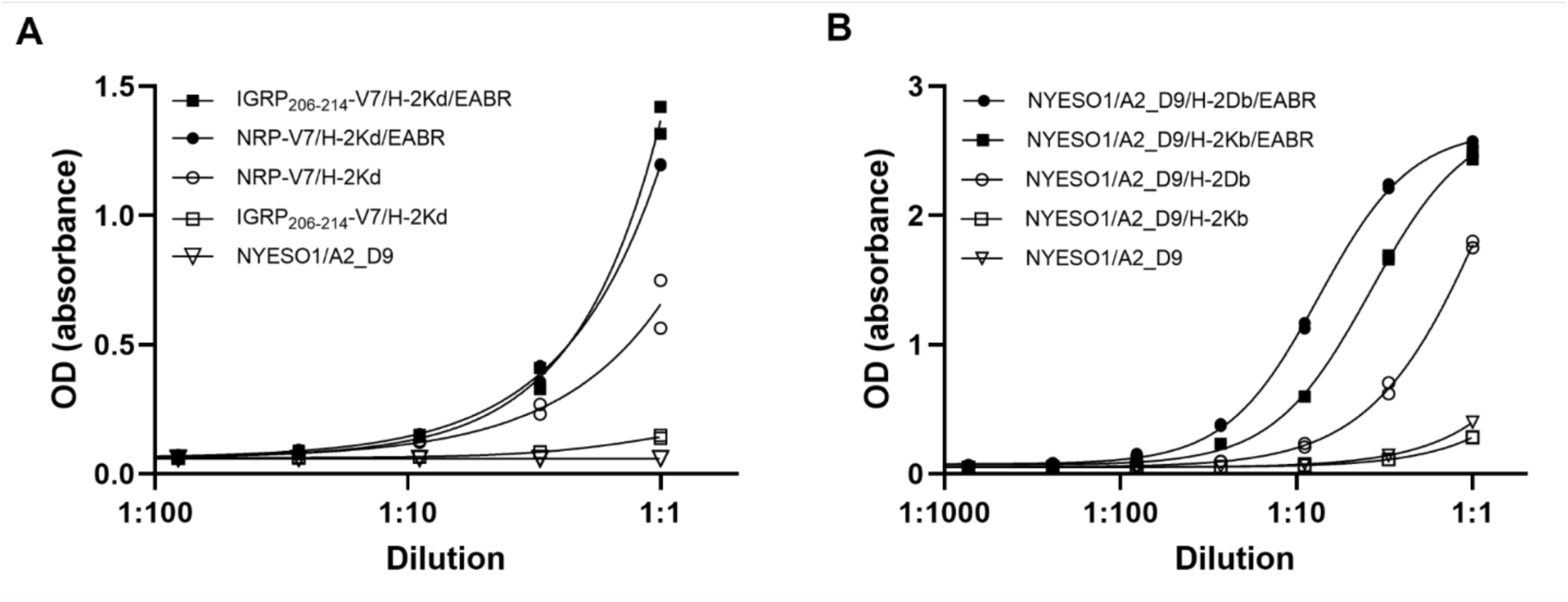
EABR-mediated APV release generalizes across SCT pMHCI variants. (A) Sandwich ELISA of murine SCT pMHCI of APVs purified by ultracentrifugation from transfected HEK293T cells (2 replicates per dilution). Mouse anti-H-2Kd capture antibody, biotinylated mouse anti-H-2Kd primary antibody, and HRP-conjugated streptavidin secondary antibody. (B) Sandwich ELISA of chimeric human/mouse SCT pMHCI protein on APVs purified by ultracentrifugation from transfected HEK293T cell culture supernatants (2 replicates per dilution). Mouse anti-NYESO1:HLA-A*02:01 capture antibody, biotinylated mouse anti-NYESO1:HLA-A*02:01 primary antibody, and HRP-conjugated streptavidin secondary antibody.

In similar fashion, the SCT form of chimeric human/mouse NYESO1/A2_D9/H-2Db pMHCI expressed substantial levels of APVs without EABR addition, whereas swapping the α3, transmembrane, and cytoplasmic domains to a different MHC haplotype, NYESO1/A2_D9/H-2Kb, led to comparatively reduced APV expression (Fig. 3B).

Mirroring the behavior of the lower-expressing murine IGRP_206-214_-V7/H-2Kd construct, addition of an EABR sequence to the cytoplasmic tail of NYESO1/A2_D9/H-2Kb increased APV expression by more than 10-fold (Fig. 3B). For both chimeric human/mouse pMHCI constructs, however, addition of the EABR sequence increased vesicle production relative to the constructs’ non-EABR counterparts.

Collectively, these results suggest that MHC haplotype and the loaded presenting peptide can influence vesicle production for SCT constructs. As shown previously by immuno-EM of the MART1/D9 and MART1/D9/EABR constructs (Fig. 2F, 2G), the increased yield of pMHCI product observed by ELISA in the non-EABR NRP-V7/H-2Kd and NYESO1/A2_D9/H-2Db constructs may be an artifact of pMHCI aggregation or indiscriminate membrane blebbing rather than intact APV formation. Regardless of this intrinsic vesicle-forming potential, addition of the EABR sequence consistently increased APV release across these different pMHCI complexes, demonstrating the generalizability of the EABR sequence for inducing APV production.

### pMHCI/EABR APVs induce IFN-γ release from their cognate T cells *in vitro*

Having confirmed that the EABR sequence reliably produces pMHCI-displaying APVs, we next asked whether EABR-induced pMHCI-displaying APVs were functional as T cell stimulators. SCT NYESO1/D9/EABR APVs were purified from transfected cell supernatants by ultracentrifugation and SEC, and the MHCI protein concentrations in purified APVs were determined by quantitative western blot analysis (Fig. 4A). An equivalent concentration of MHCI for MART1/D9/EABR, NYESO1/D9/EABR, and NYESO1:HLA-A*02:01 tetramers was mixed with donor T cells that were transduced with the 1G4 TCR specific for the NYESO1:HLA-A*02:01 epitope. T cell stimulation was evaluated using an ELISPOT assay measuring IFN-γ release (Fig. 4B).

**Figure 4.**
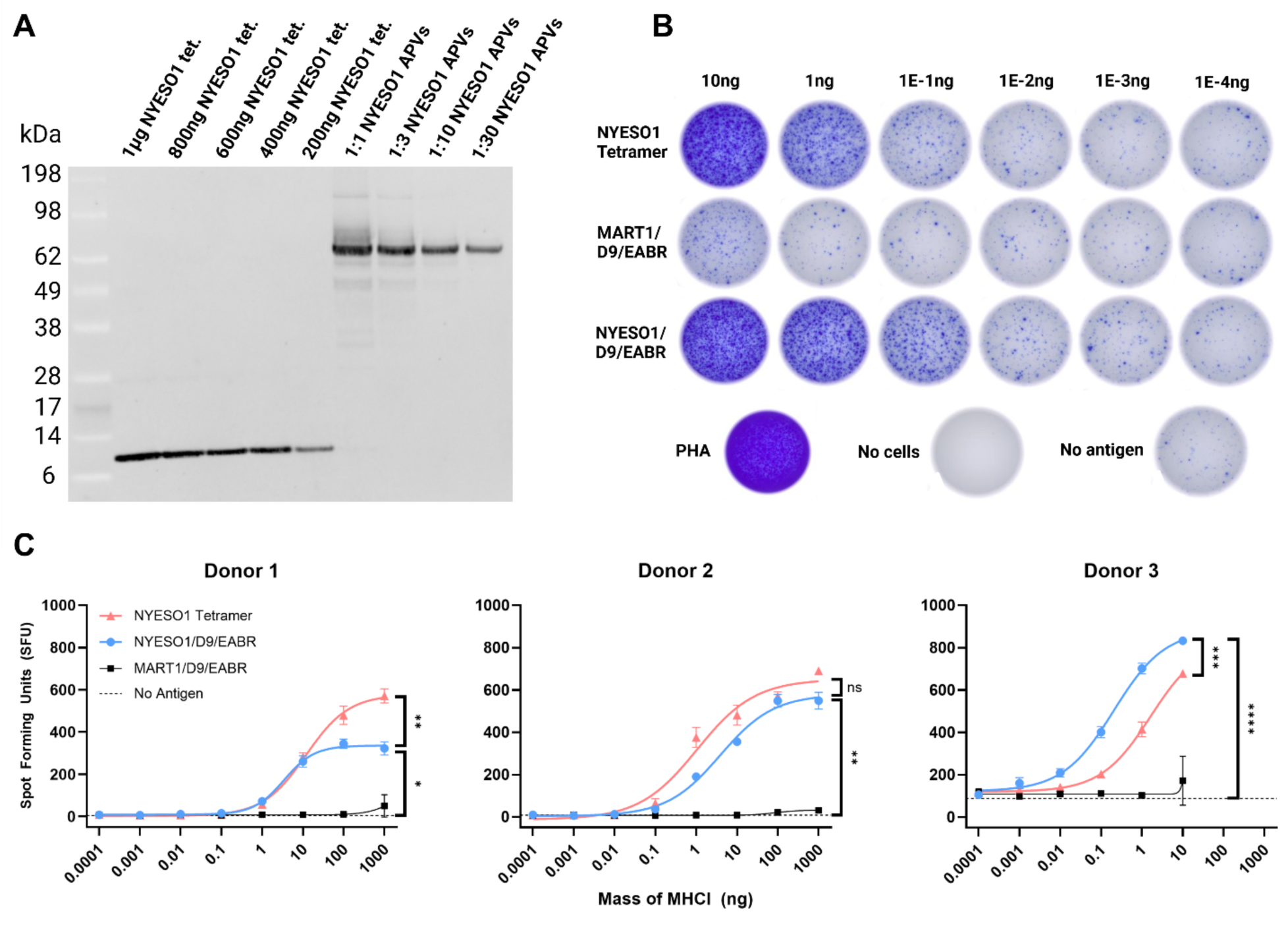
pMHCI/EABR APVs induce IFN-γ release from their cognate T cells *in vitro*. (A) Quantitative western blot comparing known amounts of NYESO1:HLA-A*02:01 tetramer (NYESO1 tet.) standards (lanes 1–5) with various dilutions of purified NYESO1/D9/EABR APVs (NYESO1 APVs) (lanes 6–9) to determine pMHCI protein concentrations in APV samples. Mouse anti-β2M primary antibody, Alexa488-conjugated anti-mouse secondary antibody. (B) Top: Scans of ELISPOT wells detecting IFN-γ release from one replicate test of Donor 3’s T cells after being dosed with serial dilutions of NYESO1:HLA-A*02:01 tetramers (NYESO1 Tetramer), SCT NYESO1/D9/EABR APVs, and SCT MART1/D9/EABR APVs. Bottom: Scans of ELISPOT wells detecting IFN-γ release from T cells dosed with 1.5% PHA as a positive control, wells without cells as a negative control (No cells), and T cells plated without antigen challenge (No antigen). (C) ELISPOT spot forming units for IFN-γ from T cells of three different donors dosed with SCT NYESO1/D9/EABR APVs, SCT MART1/D9/EABR APVs, NYESO1:HLA-A*02:01 tetramers (NYESO1 Tetramer), or no antigen challenge. Each data point is the response per 1x10^6^ cells. Baseline stimulation (i.e., “No antigen” – plated T cells without stimulation by antigen) was measured in triplicate for each donor and presented as a single mean in each plot with standard deviation. Because the NYESO1:HLA-A*02:01 tetramers generated larger spot sizes on average compared to the APVs, Donor 3’s spot forming units were too numerous to count when dosed with more than 10 ng of NYESO1:HLA-A*02:01 tetramer, preventing direct comparison between antigen conditions at higher concentrations for this donor. Significant differences between conditions linked by vertical lines are indicated by asterisks: ns > 0.05; *p < 0.05; **p < 0.01; ***p < 0.001; ****p < 0.0001.

Analysis of ELISPOT assay spot forming units (SFUs) from three different T cell donors showed that both refolded NYESO1:HLA-A*02:01 tetramers and NYESO1/D9/EABR APVs significantly stimulated 1G4 T cells above background (Fig. 4C). This level of activation was specific to the NYESO1 peptide, as 1G4 T cells dosed with MART1/D9/EABR APVs demonstrated a statistically similar response to 1G4 T cells without antigen challenge. Interestingly, the elevated baseline of T cell activation without antigen challenge in “Donor 3” coincided with an increase in the potency of NYESO1/D9/EABR APVs relative to NYESO1:HLA-A*02:01 tetramers for this donor.

The apparent association between the baseline activation level of a donor’s T cells and the potency of the NYESO1/D9/EABR APVs is also seen in the relatively weakly activated “Donor 1” and moderately activated “Donor 2” T cells, which demonstrate correspondingly weak and moderate receptiveness, respectively, to NYESO1/D9/EABR APV stimulation as compared to NYESO1:HLA-A*02:01 tetramers (Fig. S5). Future studies will explore whether this observed dynamic potency of the APVs is consistent and statistically significant.

Overall, these data confirm that the presence of pMHCI/EABR APVs is sufficient to stimulate IFN-γ release from T cells harboring their cognate TCR, and that their stimulation is specific to those T cells that feature their cognate TCR.

## Discussion

Here, we present a new approach for generating immunomodulatory pMHCI-displaying APVs from non-immune cells. SCT pMHCI/EABR constructs containing a cytoplasmic EABR sequence^22^ from the human centrosomal protein CEP55 recruit ESCRT proteins involved in cell division and viral budding to drive efficient assembly and release of pMHCI-displaying APVs. The lipid bilayer of the APV is expected to allow for pMHCI reorganization in the membrane and pMHCI clustering, both of which are considered important for the formation of a T cell stimulating immunological synapse. Purified pMHCI/EABR APVs elicited IFN-γ release from T cells, and the response was specific to T cells harboring the cognate TCR.

Unlike previous methods that have used viral coat proteins to induce budding,^32^ the pMHCI/EABR construct does not require accessory proteins for vesicle release.

Consequently, these pMHCI/EABR APVs could prove to be less immunogenic compared to other APV production methods as the pMHCI/EABR vesicles are composed of protein sequences already found in the human proteome. Thus, these T cell-targeting APVs, with putatively reduced immunogenicity, could be applicable as a targeted immunosuppressive therapeutic for autoimmune diseases. As an example, the presenting peptides that were tested for SCT murine pMHCI H-2Kd in this study – NRP-V7 and IGRP_206-214_ – are two therapeutically relevant epitopes for a murine type 1 diabetes (T1D) model; these pMHCI complexes have previously been conjugated to iron-oxide polymer nanoparticles for use as a targeted T1D therapy,^18^ suggesting a potential clinical application for the APVs described here.

Another advantage of using the EABR sequence for APV budding is that pMHCI APV assembly requires expression of only a single pMHCI/EABR protein and the corresponding gene construct could be delivered as an mRNA therapeutic. Previous work has explored the use of mRNA therapeutics for multiple sclerosis by delivering a mRNA-LNP therapeutic coding for an autogenic peptide with the goal of loading and presenting that peptide on the endogenous MHC that resides on, and is often limited to, the surface of the cell.^33^ An mRNA therapeutic coding for pMHCI/EABR may result in increased presentation of the fully formed pMHCI complex on the cell surface in addition to the release of pMHCI-loaded APVs that could distribute throughout the body, mirroring the natural release of endogenous APVs seen during an immune response and potentially increasing their therapeutic effect.^2^

The increase in purified pMHCI product resulting from the single-chain construct may be due to the single-chain heterotrimer’s increased stability.^24^ The D9 variant in particular has previously been shown to have increased traffic to the cell surface and maintains a steady-state presence there by bypassing cellular quality control steps.^27^ This increased cell surface expression and stability may be acting synergistically with the EABR sequence to increase APV budding and release in the comparatively lower-expressing HEK293T cell line, which would make the pMHCI/EABR APV approach particularly powerful as an injectable mRNA therapeutic that would likely be translated in lower-expressing primary cells in vivo. Future studies will investigate whether pMHCI/EABR APVs, or lipid nanoparticles encapsulating mRNA encoding a pMHCI/EABR construct, are effective as an autoimmune therapy.

In summary, we present a new method for efficiently generating APVs from non-immune cells. We demonstrate that these APVs display complete pMHCI complexes and are capable of specific stimulation of cognate T cells, warranting further investigation as a potential cell-free therapeutic strategy for targeted immunoregulation.

## Acknowledgements

We thank the NIH Tetramer Core Facility (contract number 75N93020D00005) for providing NYESO1:HLA-A*02:01 monomers and tetramers. We thank J. Vielmetter, A. Lam, and the Caltech Protein Expression Center for assistance with protein production; P. Gnanapragasam, A. Rorick, L. Segovia for cell culture reagents; W. Chour for MHC plasmid maps and helpful discussions. We thank D. Rees, R. Voorhees, N. Pierce, J. Keeffe, and M. Legendre for equipment access and training. We thank P. Bjorkman and O. Witte (UCLA) for helpful discussions. Immuno-electron microscopy was performed in the Caltech Cryo-EM Center with assistance from M. Ladinsky. Figures 1A, 1B, 1C, 1D, and 2C were created with Biorender.com. Research was sponsored by the U.S. Army Research Office and accomplished under cooperative agreement W911NF-19-2-0026 for the Institute for Collaborative Biotechnologies. The content of the information on this page does not necessarily reflect the position or policy of the Government, and no official endorsement should be inferred. This project has been made possible in part by grant 2023-332386 from the Chan Zuckerberg Initiative Donor Advised Fund (CZI DAF), an advised fund of the Silicon Valley Community Foundation.

## Author Contributions

B.A.O., S.L.M., and R.M.M. conceived and designed the study and analyzed the data. M.A.G.H. participated in study design. B.A.O. constructed, expressed, and evaluated DNA constructs by western blot, ELISA, DLS, Immuno-EM, and ELISPOT. K.E.H.-T. performed SEC. Z.M. created and provided 1G4 transduced T cells. B.A.O wrote the first draft of the manuscript. All authors participated in editing the manuscript.

## Materials availability

All expression plasmids generated in this study are available upon request through a Materials Transfer Agreement.

## Data and code availability

All data are available in the main text or the supplemental information. Materials are available upon request to the corresponding authors with a signed material transfer agreement. Any additional information required to reanalyze the data reported in this paper is available from the lead contact upon request. This paper does not report original code. This work is licensed under a Creative Commons Attribution 4.0 International (CC BY 4.0) license, which permits unrestricted use, distribution, and reproduction in any medium, provided the original work is properly cited. To view a copy of this license, visit https://creativecommons.org/licenses/by/4.0/. This license does not apply to figures/photos/artwork or other content included in the article that is credited to a third party; obtain authorization from the rights holder before using such material.

## Methods

### Key resources table

| REAGENT or RESOURCE | SOURCE | IDENTIFIER |
| --- | --- | --- |
| **Antibodies** |  |  |
| HLA-A Polyclonal Antibody | Thermo Fisher Scientific | Cat# PA5-29911<br>RRID: AB_2547385 |
| Recombinant Anti-HLA-A2-peptide (SLLMWITQV) Complex (3M4E5) Mouse Monoclonal Antibody | ProteoGenix | Reference:<br>RHM03002-100 |
| CD81 Monoclonal Antibody (M38) | Thermo Fisher Scientific | Cat# 10630D<br>RRID: AB_2532984 |
| CD81 Monoclonal Antibody (1.3.3.22) | Thermo Fisher Scientific | Cat# MA5-13548<br>RRID: AB_10987151 |
| CD63 Monoclonal Antibody (Ts63) | Thermo Fisher Scientific | Cat# 10628D<br>RRID: AB_2532983 |
| CD63 Polyclonal antibody | Proteintech | Cat# 25682-1-AP<br>RRID: AB_2783831 |
| Goat Anti-Human IgG Fc, Multi-Species SP ads-HRP | SouthernBiotech | Cat# 2014-05<br>RRID: AB_2795580 |
| Goat Anti-Mouse IgG Fc, Human/Bovine/Horse SP ads-HRP | SouthernBiotech | Cat# 1013-05<br>RRID: AB_2794190 |
| Goat Anti-Rabbit IgG Fab, Mouse/Human SP ads-HRP | SouthernBiotech | Cat# 4058-05<br>RRID: AB_2795982 |
| Goat anti-Rabbit IgG (H+L) Highly Cross-Adsorbed Secondary Antibody, Alexa Fluor™ Plus 488 | Thermo Fisher Scientific | Cat# A32731<br>RRID: AB_2633280 |
| Goat anti-Mouse IgG (H+L), Superclonal™ Recombinant Secondary Antibody, Alexa Fluor™ Plus 488 | Thermo Fisher Scientific | Cat# A55058<br>RRID: AB_2921066 |
| MHC Class I (H-2Kd/H-2Dd) Monoclonal Antibody (34-1-2S) | Thermo Fisher Scientific | Cat# 14-5998-82<br>RRID: AB_467822 |
| MHC Class I (H-2Kd/H-2Dd) Monoclonal Antibody (34-1-2S), Biotin | Thermo Fisher Scientific | Cat# 13-5998-82<br>RRID: AB_466872 |
| 6 nm Colloidal Gold AffiniPure™ Goat Anti-Rabbit IgG (H+L) (EM Grade) | Jackson ImmunoResearch | Cat# 111-195-144<br>RRID: AB_2338015 |
| **Bacterial and virus strains** |  |  |
| *E. coli* DH5 Alpha | Zymo Research | Cat# T3009 |
| **Chemicals, peptides, and recombinant proteins** |  |  |
| Luria Broth Base (Miller's LB Broth Base), powder | Thermo Fisher Scientific | Cat# 12795084 |
| Dulbecco's Modified Eagle Medium (DMEM) | Gibco | Cat# 11995-065 |
| Fetal bovine serum (FBS) | R&D Systems | Cat# S12450H |
| Penicillin-Streptomycin | Gibco | Cat# 15-140-122 |
| FuGENE® HD Transfection Reagent | Promega | Cat# E2311 |
| OPTI-MEM I Reduced Serum Medium | Gibco | Cat# 31985-062 |
| Expi293 Expression Medium | Gibco | Cat# A14351-01 |
| Expi293 Transfection Kit | Gibco | Cat# A14524 |
| SeeBlue Plus2 Prestained Standard | Thermo Fisher Scientific | Cat# LC5925 |
| NuPAGE™ LDS Sample Buffer | Thermo Fisher Scientific | Cat# NP0007 |
| 20X Bolt™ MES SDS Running Buffer | Thermo Fisher Scientific | Cat# B0002 |
| NucleoBond™ Xtra Maxi Plus | Macherey-Nagel | Cat# 12758412 |
| Zyppy Plasmid Miniprep Kit | Zymo Research | Cat# D4019 |
| Sucrose | Sigma-Aldrich | Cat# S5016-1KG |
| NaCl | Fisher Scientific | Cat# BP358-212 |
| Sodium hydroxide | Macron Chemicals | Cat# 7708-10 |
| Sodium bicarbonate | Macron Chemicals | Cat# 7412-12 |
| Sodium phosphate dibasic | Sigma-Aldrich | Cat# 04272-1KG |
| Tris-HCl | Fisher Scientific | Cat# BP153-1 |
| TWEEN® 20 | Sigma-Aldrich | Cat# P1379-500ML |
| BSA | Sigma-Aldrich | Cat# B4287-5G |
| Trypsin | Gibco | Cat# 25200-056 |
| L-glutamine | Gibco | Cat# 25030-149 |
| Phytohemagglutinin, M form | Gibco | Cat# 10576015 |
| Paraformaldehyde 16% Aqueous Solution EM Grade | Electron Microscopy Sciences | Cat# 15710 |
| Uranyl Formate | Electron Microscopy Sciences | Cat# 22451 |
| Biotinylated NYESO1-HLA-A*02:01 monomer | NIH Tetramer Core Facility at Emory | Task order# 73345 |
| Unlabeled NYESO1-HLA-A*02:01 tetramer | NIH Tetramer Core Facility at Emory | Task order# 73346 |
| 1-Step™ Ultra TMB-ELISA Substrate Solution | Thermo Fisher Scientific | Cat# 34029 |
| EZ-Link NHS-PEG4 Biotinylation Kit | Thermo Fisher Scientific | Cat# 21455 |
| Streptavidin (HRP) | Abcam | Cat# AB7403 |
| **Critical commercial assays** |  |  |
| Human IFN-γ Single-Color ELISPOT- Rapid | ImmunoSpot | Cat# hIFNgp-1M/5-Rapid |
| Bolt™ Bis-Tris Plus Mini Protein Gels, 4-12%, 1.0 mm, WedgeWell™ format | Thermo Fisher Scientific | Cat# NW04120BOX |
| iBlot2 NC Mini Stacks | Thermo Fisher Scientific | Cat# IB23002 |
| iBind Cards | Thermo Fisher Scientific | Cat# SLF1010 |
| iBind Flex Fluorescent Detection (FD) Solution Kit | Thermo Fisher Scientific | Cat# SLF2019 |
| iBlot2 Gel Transfer Device | Thermo Fisher Scientific | Cat# IB21001 |
| **Experimental models: Cell lines** |  |  |
| HEK293T cells | 34 | RRID:CVCL_0063 |
| Expi293F cells | Gibco | RRID:CVCL_D615 |
| **Recombinant DNA** |  |  |
| HLA-A2-peptide (SLLMWITQV) Complex (3M4E5) Human Monoclonal Antibody heavy chain | 30 | https://patentscope.wipo.int/search/en/detail.jsf?docId=EP32586798 |
| HLA-A2-peptide (SLLMWITQV) Complex (3M4E5) Human Monoclonal Antibody light chain | 30 | https://patentscope.wipo.int/search/en/detail.jsf?docId=EP32586798 |
| p3BNC-SARS-CoV-2 S-2P-DCT-EABR (a.k.a. SARSCoV2-S/EPM/EABR) | 22 | doi:10.1016/j.cell.2023.04.024 |
| D1 SCT MART1/ HLA-A*02:01 | This paper | N/A |
| D1 SCT MART1/ HLA-A*02:01/EABR | This paper | N/A |
| D9 SCT MART1/ HLA-A*02:01 | This paper | N/A |
| D9 SCT MART1/ HLA-A*02:01/EABR | This paper | N/A |
| D1 SCT NYESO1/ HLA-A*02:01 | This paper | N/A |
| D1 SCT NYESO1/HLA-A*02:01/EABR | This paper | N/A |
| D9 SCT NYESO1/ HLA-A*02:01/MHCI | This paper | N/A |
| D9 SCT NYESO1/ HLA-A*02:01/EABR | This paper | N/A |
| NYESO1/β2M | This paper | N/A |
| β2M | This paper | N/A |
| D9 SCD β2M/ HLA-A*02:01 | This paper | N/A |
| D9 SCD β2M/ HLA-A*02:01/EABR | This paper | N/A |
| D9 HLA-A*02:01 HC | This paper | N/A |
| D9 HLA-A*02:01 HC/EABR | This paper | N/A |
| NRP-V7/H-2Kd | This paper | N/A |
| IGRP<sub>206-214</sub>/H-2Kd | This paper | N/A |
| NYESO1/A2_D9/H-2Db | This paper | N/A |
| NYESO1/A2_D9/H-2Db/EABR | This paper | N/A |
| NYESO1/A2_D9/H-2Kb | This paper | N/A |
| NYESO1/A2_D9/H-2Kb/EABR | This paper | N/A |
| **Software and algorithms** |  |  |
| Image J 1.54f | NIH | https://imagej.net/<br>RRID: SCR_003070 |
| Prism 10.3.1 | GraphPad |  |
| **Other** |  |  |
| Amicon Ultra-15 100 kDa MWCO Centrifugal Filter Unit | EMD Millipore | Cat# UFC910096 |
| Amicon Ultra-4 10 kDa MWCO Centrifugal Filter Unit | EMD Millipore | Cat# UFC801024 |
| Nanosep with 100K Omega | Pall Life Sciences | Cat# OD100C34 |
| Erlenmeyer cell culture flasks - 500mL | Corning | Cat# 431145 |
| 6-well Clear Flat Bottom TC-treated Multiwell Cell Culture Plate, with Lid | Falcon | Cat# 353046 |
| 30 mL BD Luer-Lok™ Syringe | BD | Cat# 302832 |
| 3 mL BD Luer-Lok™ Syringe | BD | Cat# 309657 |
| AirOtop enhanced seal for 125 ml ultra yield flask | VWR | Cat# 899421 |
| Costar® Spin-X® Centrifuge Tube Filters, 0.22 μm Pore CA Membrane | Corning | Cat# 8160 |
| Polycarbonate centrifuge tubes | Beckman Coulter | Cat# 349622 |
| 0.45um syringe filter | Corning | Cat# 431220 |
| 0.2um syringe Filter | VWR | Cat# 28143-300 |
| Nalgene™ Rapid-Flow™ Sterile Disposable Filter Unit - 0.45um | Thermo Fisher Scientific | Cat# 156-4045 |
| Costar high binding plates, 96-well | Corning | Cat# 9018 |
| TempPlate Sealing Foil | USA Scientific | Cat# 2923-0100 |
| HiTrap MabSelect | Cytiva | Cat# 28408253 |
| Superose 6 Increase 10/300 column | Cytiva | Cat# 29-0915-96 |
| HiLoad 16/600 Superdex 200 column | Cytiva | Cat# 28-9893-35 |
| Formvar/Carbon 300 Mesh grids | Electron Microscopy Sciences | Cat# FCF300-Cu-50 |

## Experimental Model and Subject Details

### Bacteria

*E. coli* DH5 Alpha cells (Zymo Research) used for expression plasmid productions were cultured in LB broth (Sigma-Aldrich) with shaking at 225 rpm at 37 °C. Plasmids were purified for transfection using Zymogen miniprep kits (Zymo Research) or maxiprep kits (Macherey-Nagel).

### Cell lines

HEK293T cells plated in 6-well plates (Falcon) were cultured in Dulbecco’s modified Eagle’s medium (DMEM, Gibco) supplemented with 10% heat-inactivated fetal bovine serum (FBS, R&D Systems) and 1 U/mL penicillin-streptomycin (Gibco) at 37 °C and 5% CO_2_. Transfections were carried out with FuGENE transfection reagent (Promega) diluted in Opti-MEM (Gibco). Expi293F cells (Gibco) for APV expression and antibody expression were maintained at 37 °C and 8% CO_2_ in Expi293 expression medium (Gibco). Transfections were carried out with an Expi293 Expression System Kit (Gibco). Falcon tubes sealed with AirOtops (VWR) or culture flasks (Corning) containing Expi293F cells were maintained under shaking at 470 rpm for 10-30 mL transfections and 130 rpm for transfections larger than 30 mL. All cell lines were derived from female donors and were not specially authenticated.

### T cells

T cells were isolated from human donor whole blood samples by negative selection using the RosetteSep Human T Cell Enrichment Cocktail Kit (STEMCELL Technologies). Isolated T cells were seeded in a fresh complete ImmunoCult™-XF T cell expansion medium (STEMCELL Technologies) at 1 x 10^6^ cells/mL with 2 μL/mL ImmunoCult™ human CD3/CD28 T cell activator. T cells were activated for 3 days and expanded for up to 12 days by changing into fresh expansion medium every 2-3 days. All cells were cultured in incubators at 37 °C and 5% CO_2_.

## Methods

### Design of EABR constructs

The EABR sequence was identical to the sequence used by Hoffmann et al. to create SARS-CoV2-S eVLPs.^24^ Briefly, the EABR domain (residues 160-217) of the human CEP55 protein was fused to the C-terminus of the SCT MART1/HLA-A*02:01 alpha chain separated by a (Gly)_3_Ser (GS) linker to generate “D1” and “D9” single-chain heterotrimer pMHCI constructs inserted in the p3BNC expression plasmid. To generate the “D1” SCT construct, the N-terminus of the HLA-A*02:01 alpha chain was extended with a 4x(G_4_S) linker connecting to the C-terminus of β2M, which was itself extended at its N-terminus by a 3x(G_4_S) linker connecting to either the 10-mer MART1 (ELAGIGILTV) or 9-mer NYESO1 (SLLMWITQV) cancer-related peptides to make a single-chain heterotrimer (SCT)^22^ (Fig. 1B). To generate the SCT “D9” construct, the “D1” SCT construct was edited with Y84C and A139C substitutions.^26^

The “single-chain dimer” (SCD) construct removed both the peptide and the 3x(G_4_S) linker between the peptide and β2M of the “D9” SCT construct. The “heavy chain” (HC) construct removed the peptide, the 3x(G_4_S) linker between the peptide and β2M, β2M, and the 4x(G_4_S) linker between β2M and the MHCI heavy chain of the “D9” SCT construct (Fig. 2C). Both the SCD and HC constructs were inserted in p3BNC expression plasmids.

Murine SCT MHCI H-2Kd was constructed similar to the human D1 SCT HLA-A*02:01 design, with either an NRP-V7 or IGRP_206-214_ presenting peptide fused to a 3x(Gly_4_Ser) linker, murine β2M, a 4x(Gly_4_Ser) linker, and the H-2Kd heavy chain, i.e., NRP-V7/H-2Kd and IGRP_206-214_/H-2Kd, respectively. The chimeric human/mouse MHCI HLA-A2/H-2Db and HLA-A2/H-2Kb were designed according to the D9 SCT design, with an NYESO1 presenting peptide fused to a 3x(Gly_4_Ser) linker, human β2M, a 4x(Gly_4_Ser) linker, and a chimeric MHCI alpha chain composed of the extracellular α1 and α2 domains of human Y84A,A139C HLA-A*02:01 heavy chain and the α3, transmembrane, and cytoplasmic domains of murine H-2Kb or H-2Db heavy chain, i.e., NYESO1/A2_D9/H-2Db and NYESO1/A2_D9/H-2Kb, respectively. Murine and chimeric human/murine SCT pMHCI constructs were inserted into the p3BNC expression plasmid.

### Production of EABR APVs

EABR APVs were generated by one of two means: 1) transfecting Expi293F cells (Gibco) cultured in Expi293F expression media (Gibco) on an orbital shaker at 37 °C and 8% CO2 with plasmid DNA pre-filtered through 0.22 µm Spin-X filters (Corning); 2) transfecting HEK293T cells cultured in Dulbecco’s modified Eagle’s medium (DMEM, Gibco) supplemented with 10% heat-inactivated fetal bovine serum (FBS, Sigma-Aldrich) and 1 U/ml penicillin-streptomycin (Gibco) at 37 °C and 5% CO_2_. 72 hours post-transfection, cells were centrifuged at 1000 x g for 10 min, supernatants were passed through a 0.45 um filter (Corning) with Luer-Lok syringes (BD) or a 0.45 um vacuum filter (Thermo Fisher Scientific), and concentrated using Amicon Ultra-15 centrifugal filters with 100 kDa molecular weight cut-off (Millipore). APVs were purified by ultracentrifugation at 50,000 rpm (135,000 x g) for 2 hours at 4 °C using a TLA100.3 rotor and an OptimaTM TLX ultracentrifuge (Beckman Coulter) on a 20% w/v sucrose cushion in polycarbonate centrifuge tubes (Beckman Coulter). Supernatants and the sucrose cushion were removed, and pellets were re-suspended in 200 uL sterile pH 7.4 PBS at 4 °C overnight. To remove residual cell debris, re-suspended samples were transferred to microcentrifuge tubes and centrifuged at 10,000 x g for 10 min; clarified supernatants were collected for subsequent ELISA, DLS, and western blot assays. For immuno-EM grid preparations, quantitative western blots, and ELISPOT assays, APVs were further purified by SEC using a Superose 6 Increase 10/300 column (Cytiva) equilibrated with pH 7.4 PBS. Peak fractions in the void volume corresponding to pMHCI APVs were combined and concentrated to 250-500 uL in Amicon Ultra-4 centrifugal filters with 100 kDa molecular weight cut-off (Millipore) or Nanosep with 100K Omega centrifugal filters (Pall). Samples were stored at 4 °C and imaged directly after purification.

### Transduction of TCRs in PBMC

Engineering of candidate TCRs in PBMC was performed according to previous publication.^35^

### Antibody Expression

Anti-HLA-A2-peptide (SLLMWITQV) Complex (3M4E5) antibody was made in-house by expressing the heavy chain and light chain of the 3M4E5 antibody at a 1.5:1 plasmid ratio in Expi293F cells.^33^ Supernatant from the transiently-transfected Expi293F cells (Gibco) was separated from the cells by pelleting at 3500g for 15 minutes and filtering the clarified supernatant through a 0.2 um syringe filter (VWR) or 0.45 um vacuum filter unit (Thermo Fisher Scientific). The filtered supernatant was further purified using Fc-affinity chromatography (HiTrap MabSelect, Cytiva) and SEC (HiLoad 16/600 Superdex 200 column, Cytiva). Peak fractions corresponding to purified antibody proteins were pooled, concentrated, and stored at 4 °C. Biotinylated antibodies for ELISAs were generated and purified using an EZ-Link biotinylation kit (Thermo Fisher Scientific). The in-house generated antibody was compared to a commercially available 3M4E5 antibody (RHM03002-100; ProteoGenix) with ELISA and showed identical performance (data available upon request).

### Western blot analysis

The presence of HLA-A, C81, and CD63 on purified APVs was detected by Western blot analysis. Samples were diluted in NuPage SDS-PAGE loading buffer (Thermo Fisher Scientific) under reducing conditions, separated on Bolt™ Bis-Tris Plus Mini Protein Gels, 4-12% (Thermo Fisher Scientific), and transferred to nitrocellulose membranes in the form of an iBlot2 NC Mini Stack (Thermo Fisher Scientific) using the iBlot2 Gel Transfer Device (Thermo Fisher Scientific). Nitrocellulose membranes from the stack were transferred to water for 1-5 minutes, then transferred to Flex FD solution (Thermo Fisher Scientific), before placing on an iBind card (Thermo Fisher Scientific) for staining with antibodies per the iBlot2 protocol. The following antibodies were used for detecting pMHCI: rabbit anti-HLA-A protein (PA5-29911; Thermo Fisher Scientific) at 1:1,000, mouse anti-CD81 (10630D; Thermo Fisher Scientific) at 1:100, mouse anti-CD81 (MA5-13548; Thermo Fisher Scientific) at 1:100, mouse anti-CD63 (10628D; Thermo Fisher Scientific) at 1:500, rabbit anti-CD63 (25682-1-AP; Thermo Fisher Scientific) at 1:500, Alex488-conjugated goat anti-rabbit IgG (A32731; Thermo Fisher Scientific) at 1:1,000, and Alex488-conjugated goat anti-mouse IgG (A55058; Thermo Fisher Scientific) at 1:1,000. Protein was compared to a SeeBlue Plus2 Prestained protein ladder (Thermo Fisher Scientific). For T cell studies, the amount of MHCI on APVs was determined by quantitative Western blot analysis. Various dilutions of SEC-purified pMHCI APV samples and known amounts of soluble, biotinylated pMHCI protein (NIH Tetramer Core Facility at Emory) were separated by SDS-PAGE and transferred to nitrocellulose membranes as described above. Band intensities of the pMHCI standards and pMHCI APV sample dilutions were measured using ImageJ to determine MHCI concentrations.

### Immuno-EM of pMHCI/EABR APVs

SEC-purified MART1/MHCI/EABR APVs were prepared on grids for negative stain transmission electron microscopy at room temperature. Formvar/Carbon 300 Mesh grids (Electron Microscopy Sciences) were first glow discharged. 20 uL of purified sample was pipetted onto paraffin, glow discharged formvar/carbon 300 mesh grids were placed on the droplet of sample for 5 minutes, then gently wicked away by dabbing the edge of the grid on filter paper. The grids were then placed on a 20uL droplet of 1% paraformaldehyde (Electron Microscopy Sciences) in pH 7.4 PBS for 10 minutes, and subsequently wicked away. Grids were washed three times by placing the grids on droplets of pH 7.4 PBS for 5 minutes, and wicking away the PBS with filter paper after each wash. Grids were then blocked by placing the grids on droplets of 5% FBS diluted in pH 7.4 PBS for one hour. Blocking solution was wicked away from the grids, and the grids were then placed on droplets of rabbit anti-HLA-A antibody (PA5-29911; Thermo Fisher Scientific) diluted 1:100 dilution in 5% sucrose, 5% FBS pH 7.4 PBS for 2 hours; to prevent drying of the primary antibody solution, the grids were placed in a humidified chamber made from wet paper towels folded under a pyrex glass dish. Grids were again washed three times by placing the grids on droplets of pH 7.4 PBS for 10 minutes, and wicking away the PBS with filter paper after each wash. The grids were then placed on droplets of 6 nm colloidal gold goat anti-rabbit antibody (PA5-29911; Thermo Fisher Scientific) diluted 1:20 dilution in 5% sucrose, 5% FBS pH 7.4 PBS for 1 hour. Grids were then washed three times, ten minutes per wash with pH 7.4 PBS, wicking away the PBS after the ten minute wash each time. Finally, the grid was placed on a droplet of 1.5% uranyl formate (Electron Microscopy Sciences), prepared fresh within 1 week of use, for 1 minute. The excess uranyl formate was wicked away and the grid was left to air dry for 30 minutes before storing at room temperature away from light. Imaging of the grids was performed on a FEI Tecnai T12 Transmission Electron Microscope within two weeks after the grids were prepared.

### Indirect ELISA

96-well plates (Corning) were coated with APV samples diluted 1:10 or 1:4 or 1:1, 4-fold serially diluted in sterile 0.1 M NaHCO_3_ pH 9.6, and sealed with TempPlate sealing foil (USA Scientific) overnight at 4 °C. Plates were emptied and blocked with TBS-T/3% BSA (Sigma-Aldrich) for at least 30 minutes. Plates were again emptied, and 5 ug/mL of primary antibody diluted in TBS-T/3% BSA was added to the plates. The following were used as primary antibodies for detecting pMHCI: rabbit anti-HLA-A antibody (PA5-29911; Thermo Fisher Scientific), mouse anti-HLA-A2-peptide (SLLMWITQV) Complex (3M4E5) antibody (RHM03002-100; ProteoGenix), mouse anti-H-2Kd antibody (14-5998-82; Thermo Fisher Scientific).

After a 2-hour incubation at room temperature, plates were emptied, washed three times with TBS-T, and 5 ug/mL of HRP-conjugated secondary antibody diluted in TBS-T/3% BSA was added to the plates for 30 minutes at room temperature. The following were used as secondary antibodies: HRP-conjugated goat anti-rabbit IgG (4058-05; Southern Biotech) at 1:2,000, HRP-conjugated goat anti-mouse IgG (1013-05; Southern Biotech) at 1:2,000.

After washing three times with TBS-T, plates were developed using 1-Step™ Ultra TMB-ELISA Substrate Solution (Thermo Fisher Scientific) and absorbance was measured at 450 nm. Standard 4-parameter sigmoidal binding curves were calculated using GraphPad Prism 10.3.1 without any further editing.

### Sandwich ELISA

96-well plates (Corning) were coated with 5 ug/mL of mouse anti-HLA-A2-peptide (SLLMWITQV) Complex (3M4E5) antibody (RHM03002-100; ProteoGenix) or in-house-made anti-HLA-A2-peptide (SLLMWITQV) Complex (3M4E5) antibody diluted in sterile 0.1 M NaHCO_3_ pH 9.6, and sealed with TempPlate sealing foil (USA Scientific) overnight at 4 °C. Plates were emptied and blocked with TBS-T/3% BSA for at least 30 minutes. Plates were again emptied, and APV samples diluted 1:10 or 1:4, 4-fold serially diluted in TBS-T/3% BSA were added to the plates. After a 2-hour incubation at room temperature, plates were emptied, and 5 ug/mL of biotinylated detection antibody diluted in TBS-T/3% BSA was added to the plates. 5 ug/mL of biotinylated (Thermo Fisher Scientific) mouse anti-HLA-A2-peptide (SLLMWITQV) Complex (3M4E5) antibody (RHM03002-100; ProteoGenix) or in-house-generated human anti-HLA-A2-peptide (SLLMWITQV) Complex (3M4E5) antibody was used as a capture antibody. After another 2-hour incubation at room temperature, plates were washed three times with TBS-T. HRP-conjugated streptavidin (Abcam) was diluted to manufacturer’s recommendations in TBS-T/3% BSA and added to plates for 30 minutes at room temperature. After washing three times with TBS-T, plates were developed using 1-Step™ Ultra TMB-ELISA Substrate Solution (Thermo Fisher Scientific) and absorbance was measured at 450 nm. Standard 4-parameter sigmoidal binding curves were calculated using GraphPad Prism 10.3.1 without any further editing.

### ELISPOT

1G4 T cells were centrifuged at 300g for 10 min, and resuspended in CTL-TestTM media (ImmunoSpot) containing 1% L-glutamine (Gibco) for ELISPOT analysis to evaluate T cell responses. APVs were added to human IFN-g single-color ELISPOT plates (ImmunoSpot) and 10-fold serially diluted. Phytohemagglutinin (Gibco) was added in a separate well at a final concentration of 1.5% as a positive control. 100,000 cells were added per well, and plates were incubated at 37 °C in 8% CO_2_ for 24 hours. Biotinylated detection, streptavidin-alkaline phosphatase (AP), and substrate solutions were added according to the manufacturer’s guidelines. Plates were gently rinsed with water three times to stop the reactions. Plates were air-dried for two hours in a running laminar flow hood. The number of spots and the mean spot sizes were quantified using a CTL ImmunoSpot S6 Universal-V Analyzer (Immunospot).

### Quantification and Statistical Analysis

ELISA serial dilutions were plotted by fitting a standard sigmoidal four parameter logistic curve without weighting or special handling of outliers using Graphpad Prism 10.3.1. Statistical significance in ELISPOT data was calculated using multiple unpaired t-tests with Welch correction and Holm-Šídák’s multiple comparison tests to obtain adjusted p-values in Graphpad Prism 10.3.1; unless otherwise stated, adjusted p-values are reported for comparisons of the maximum concentrations.

---

**Figure S1.**
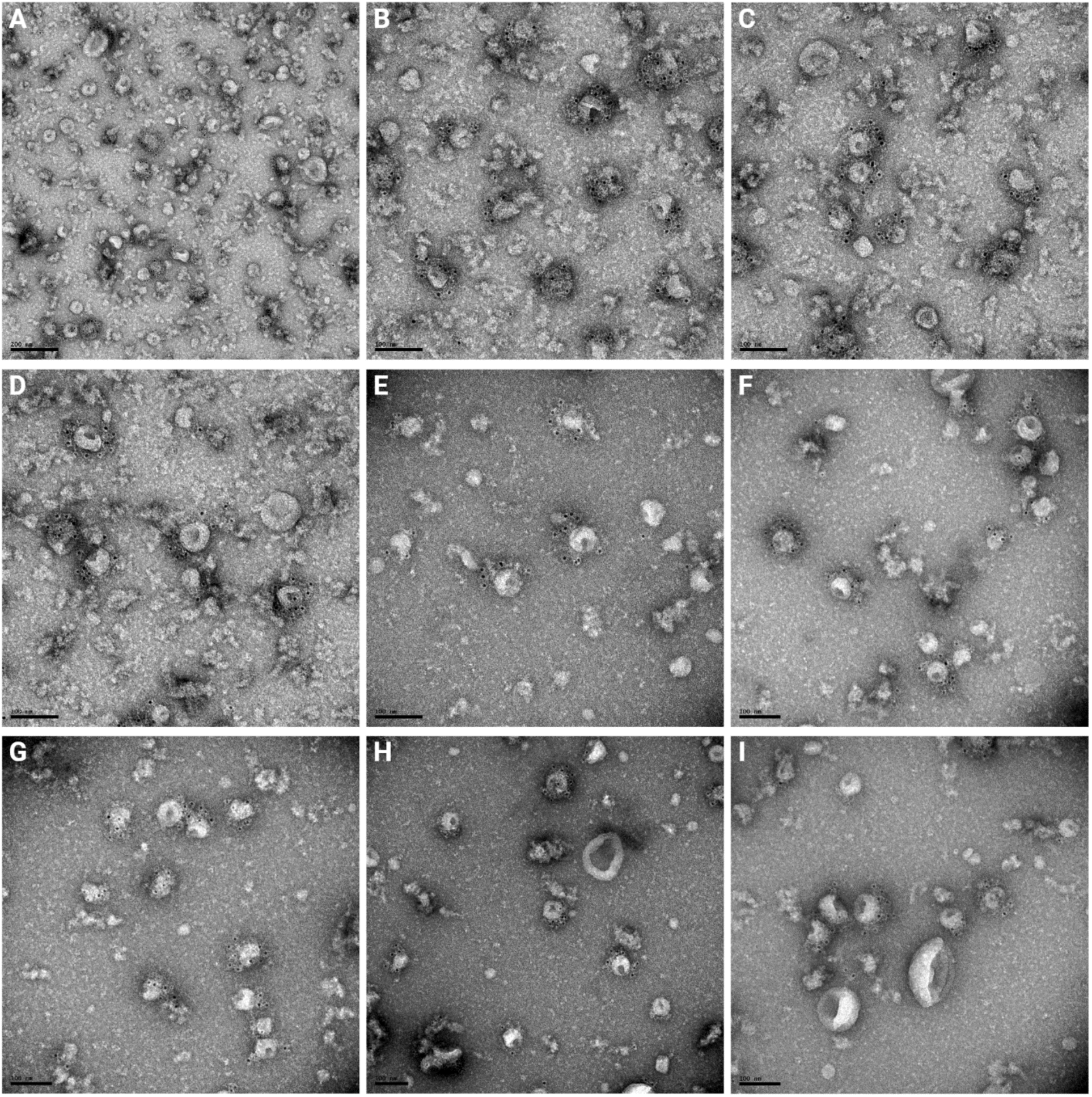
Additional micrographs of pMHCI/EABR grids prepared by immuno-EM show APVs. All micrographs show Expi293F cell culture supernatant after transfection with a SCT MART1/D1/EABR construct and purified by ultracentrifugation and SEC as outlined in Figure 1C. (A-D): 50 mL of purified Expi293F cell culture supernatant. (E-I) 200 mL of purified Expi293F cell culture supernatant further diluted in pH 7.4 PBS after purification. All grids were prepared using the identical protocol as in Figures 1F and 2D-F. Secondary 6 nm gold-conjugated anti-rabbit antibody bound to primary rabbit anti-HLA-A antibody appears as black punctate in the image. (A) Scale bar, 200 nm. (B-I) Scale bar, 100 nm.

**Figure S2.**
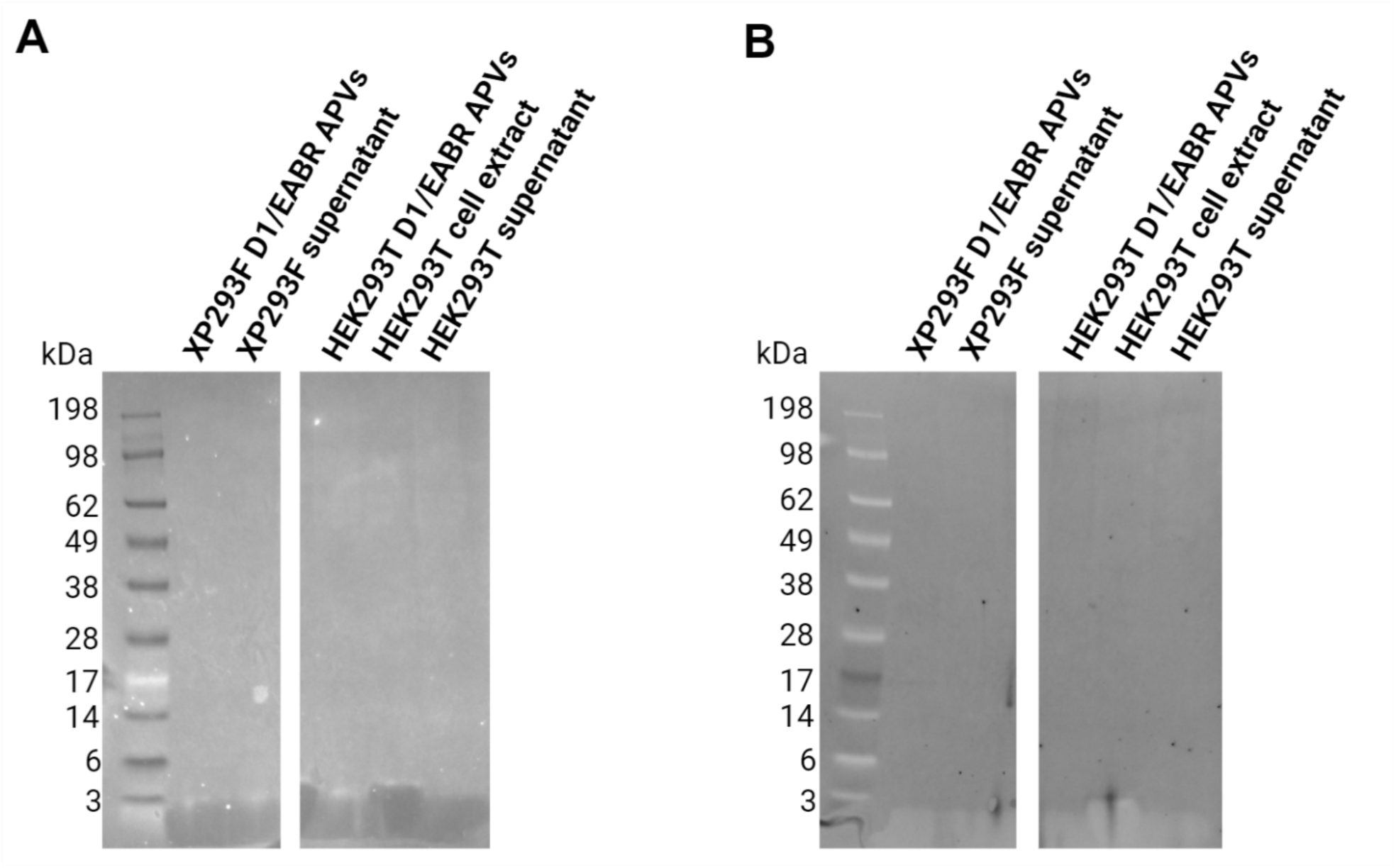
Western Blots of purified APVs are negative for CD81 and CD63. (A-B) Western Blots of purified Expi293F SCT D1 MART1/D1/EABR APVs (XP293F D1/EABR APVs), Expi293F purified supernatant, HEK293T SCT D1 MART1/D1/EABR APVs (HEK293T D1/EABR APVs), HEK293T cell extract, and HEK293T purified supernatant. (A) Western Blot of CD81. (B) Western Blot of CD63.

**Figure S3.**
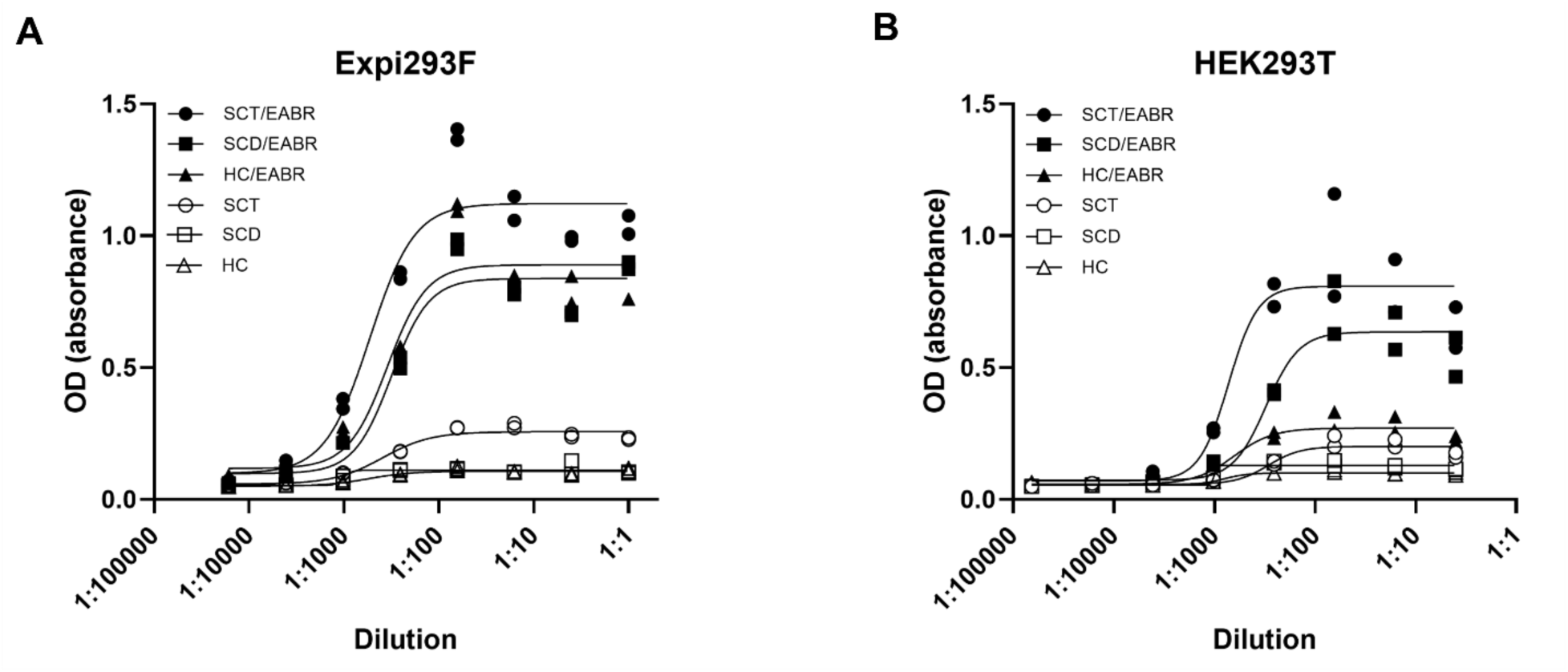
Indirect ELISA of purified Expi293F and HEK293T transfection supernatant suggests complete pMHCI not required for APV release. Indirect ELISA of purified Expi293F (A) and HEK293T (B) supernatant transfected with SCT MART1/D1/EABR (SCT/EABR), SCD D1/EABR (SCD/EABR), D1 HC-EABR (HC/EABR), and the non-EABR versions of the three constructs just described: SCT MART1/D1 (SCT), SCD D1 (SCD), and HC D1 (HC). See Figure 2C for diagrams of SCT, SCD, and HC. All supernatant was purified by ultracentrifugation and resuspended in pH 7.4 PBS. Polyclonal rabbit anti-HLA-A was used for the primary antibody, and HRP-conjugated goat anti-rabbit was used for the secondary antibody. (2 replicate measurements per dilution).

**Figure S4.**
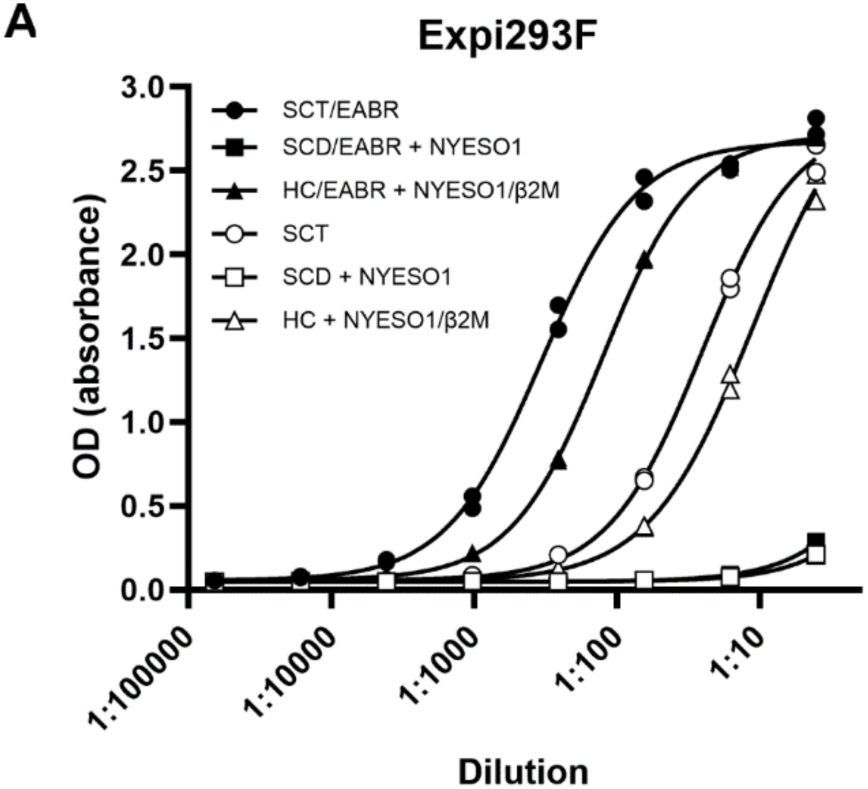
Sandwich ELISA of purified Expi293F transfection supernatant shows non-EABR SCT capable of APV formation. Sandwich ELISA of purified Expi293F supernatant transfected with: SCT NYESO1/D9/EABR (SCT/EABR); SCD D9/EABR (SCD/EABR) along with co-transfected NYESO1 peptide; HC-D9/EABR (HC/EABR) along with co-transfected NYESO1/β2M; and, the non-EABR versions of the three constructs just described.— SCT NYESO1/D9 (SCT), SCD D9 (SCD) with co-transfected NYESO1, and HC-D9 (HC) with co-transfected NYESO1/β2M. All supernatant was purified by ultracentrifugation and resuspended in pH 7.4 PBS. Monoclonal human anti-NYESO1:HLA-A*02:01 complex was used for the initial capture antibody. Biotinylated monoclonal human anti-NYESO1:HLA-A*02:01 complex was used for the primary detection antibody and secondary strep-HRP was then incubated in each well before developing with HRP substrate. (2 replicate measurements per dilution).

**Figure S5.**
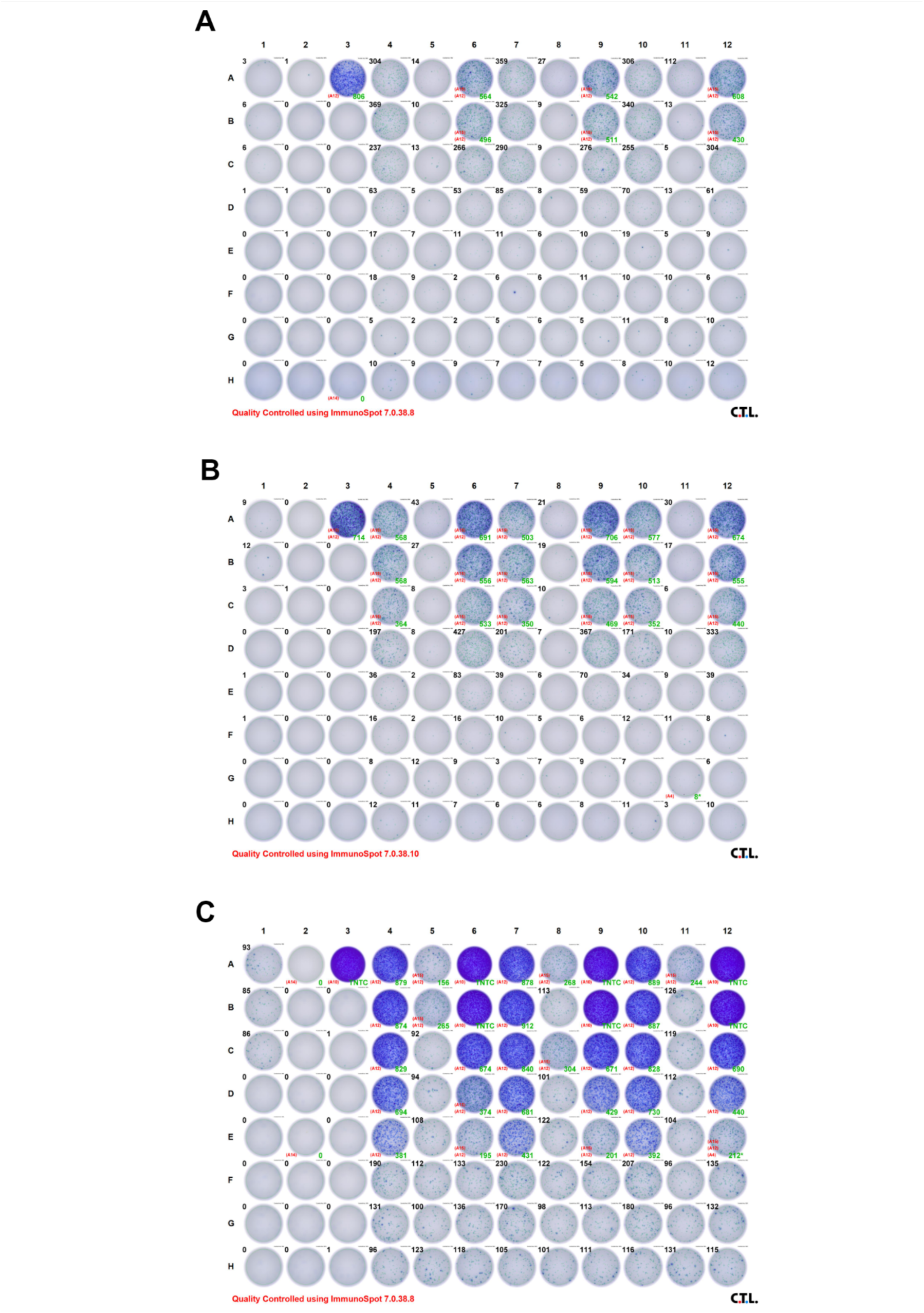
Scans of ELISPOT plates detecting IFN-γ release from donor T cells after APV co-incubation. (A-C) Scans of ELISPOT wells detecting IFN-γ release from donor T cells after being dosed with serial dilutions of SCT NY-ESO-1/D9/EABR APVs (lanes 4,7,10), SCT MART1/D9/EABR APVs (lanes 5,8,11), and NYESO1:HLA-A2 tetramers (lanes 6,9,12); 1.5% PHA (lane 3, row A); without cells or antigen challenge (lane 2, rows A); and cells without antigen challenge (lane 1, rows A-C). (A) Donor 1; (B) Donor 2; (C) Donor 3.

